# Accuracy-speed-stability trade-offs in a targeted stepping task are similar in young and older adults

**DOI:** 10.1101/2022.12.15.520631

**Authors:** Wouter Muijres, Sylvie Arnalsteen, Cas Daenens, Maarten Afschrift, Friedl De Groote

## Abstract

**Background:** Stepping accuracy, speed, and stability are lower in older compared to young adults. Lower stepping performance in older adults may be due to larger accuracy-speed-stability trade-offs because of reduced ability to simultaneously fulfill these task-level goals. Our goal was to evaluate whether trade-offs are larger in older compared to young adults in a targeted stepping task. Since sensorimotor function declines with age, our secondary goal was to evaluate whether poorer sensorimotor function was associated with larger trade-offs.

**Methods:** 25 young (median 22 years old) and 25 older (median 70 years old) adults stepped into projected targets in conditions with various levels of accuracy, speed, and stability requirements. We determined trade-offs as the change in performance, i.e. foot placement error, step duration, and mediolateral center of pressure path length, between each of these conditions and a control condition. To assess age-related differences in the magnitude of trade-offs, we compared the change in performance between age groups. Associations between trade-offs and measures of sensorimotor function were tested using correlations.

**Results:** We found an accuracy-speed and an accuracy-stability trade-off in both young and older adults, but trade-offs were not different between young and older adults. Inter-subject differences in sensorimotor function could not explain inter-subject differences in trade-offs.

**Conclusion:** Age-related differences in the ability to combine task-level goals do not explain why older adults stepped less accurate and less stable than young adults. However, lower stability combined with an age-independent accuracy-stability trade-off could explain lower accuracy in older adults.

## Introduction

One out of three adults aged over 65 years old falls at least once a year (1–3). Falls have been reported to result in injuries (33% minor and 15% serious) and even deaths (1,4). The most common cause of falling is incorrect body weight shifting (5). Weight shifts are of great importance to locomotor tasks such as stepping as adequate weight shifts are a prerequisite to lift the swing foot of the ground (6,7). In young adults, pre-step weight shifts have been found to determine stepping accuracy and speed (7,8). Incorrect weight shifts may thus challenge balance by reducing the ability for timely and accurate foot placement to maintain balance. When the task is to step into a target, stepping accuracy may come at the cost of stability as restraining foot placement reduces available balance strategies to non-stepping strategies (9,10). In addition, there is a trade-off between speed and accuracy. Healthy young adults step slower when they are required to step more accurate (11,12). Compared to young adults, older adults step slower (8,13–15) and less accurate (16–18). It is yet unclear whether reduced stepping speed and accuracy in older adults are related to balance control problems. Older adults may have more trouble than young adults to combine different stepping goals resulting in a larger decrease in performance on one task-level goal (e.g. accuracy) when the demands on another task-level goal are increased (e.g. speed or stability). The primary aim of our pre-registered study was to determine whether trade-offs between stepping speed, accuracy, and stability are larger in older compared to young adults.

Stepping accuracy, speed, and stability are reduced in older compared to young adults. Older adults step less accurate to targets during walking (16,17). This is particularly true when the time to respond to an appearing target is short (17). In gait initiation, older adults position their foot less accurate when stepping towards targets that move laterally and late during step initiation (18). In addition to lower accuracy, older adults are slower in forward stepping (8,15) and in a choice stepping task, where the target could appear in multiple positions (8,13,14). Stepping stability as derived from center of pressure/mass kinematics, is lower in older than in young adults. More specifically, pre-step mediolateral center of pressure fluctuations and end-of-step mediolateral center of mass displacement are higher in older compared to young adults (15,19). However, to our knowledge, age-related changes in stability have not been associated with speed or accuracy in targeted stepping.

Decreased stepping accuracy, speed, and stability in older adults may result from the inability to simultaneously fulfill these three task level goals. The trade-off between speed and accuracy has been well documented. It was initially studied in upper limb reaching tasks but the same trade-off has been found when stepping to a target (11,20,21). In young adults, foot positioning was slower when demands on accuracy increased, such as smaller targets or larger movement amplitudes (11). The other way around, foot placement accuracy decreased with increasing speed in targeted stepping (20). However, it is unclear whether there is an effect of age on the speed-accuracy trade-off. The accuracy-stability trade-off has been studied less extensively (22,23). External balance support, that reduces balance control requirements during a stepping task, improves stepping accuracy (22,23). It is however unclear if aging influences the accuracy-stability trade-off. The existance of a speed-stability trade-off in older adults has been suggested but has not been tested. For example, Luchies et al. (8) and Patla et al. (13) observed that older adults performed slower postural movements and larger weight shifts in preparation of a step and interpreted this as a strategy to increase stability at the cost of speed. Here, we systematically studied the effect of age on the trade-offs between speed, accuracy, and stability by manipulating stepping conditions.

Stepping accuracy, speed, and stability trade-offs may be larger in older than in young adults due to sensorimotor decline. Sensorimotor function declines with age resulting in higher motor noise (24) and reduced sensory function (25–27). The speed-accuracy trade-off has been suggested to emerge from motor noise (28) therefore age-related increases in motor noise may induce a larger speed-accuracy trade-off. Motor noise is signal dependent and therefore fast movements, requiring higher muscle activations and stronger muscle contraction, may result in more variable force production and thereby lower accuracy. As sensorimotor function is decreased in older adults, this effect may be larger in older compared to young adults. In addition, sensorimotor decline has been repeatedly associated with balance deficits (29–33). Indeed, results from our lab suggest that increased sensorimotor noise can explain age-related alterations in reactive standing balance, such as increased peak hip angle as a response to a support surface translation (34). Age-related decreases in sensorimotor function may thus affect the ability to meet accuracy, speed, and stability goals simultaneously. The secondary aim of this study was therefore to explore the relationship between age-related decline in sensorimotor function and task-level goal trade-offs.

Summarized, our confirmatory hypothesis was that accuracy-speed-stability trade-offs would be larger in older compared to young adults. Moreover, our exploratory hypothesis was that larger accuracy-speed-stability trade-offs would be associated with poorer sensorimotor function. To test these hypotheses, we studied stepping into a target in young and older adults in a range of conditions with different accuracy, speed, and stability demands. For each condition, we measured accuracy, speed, and stability performance. We defined task-level goal trade-offs as the change in stepping performance for one task level goal (e.g. speed) as an effect of increasing the demands for the other task level goals (e.g. accuracy or stability). We tested for differences in trade-offs between groups. Additionally, we quantified sensorimotor function and performed a correlation analysis to test relationships between sensorimotor outcomes and task-level goal trade-offs.

## Methods and protocol

### Subjects

Twenty-five healthy young (median = 22, range = [20, 29] years, 12 females) and 25 healthy older adults (median = 70, range = [67, 89] years, 13 females) participated in the study, see Table 1. Prior to participation, subjects completed the Fall Efficacy Scale-International (FES-I) and the MoCA (Montreal Cognitive Assessment). Subjects were excluded when they were suffering from severe obesity (BMI > 30), severe orthostatic hypotension, acute neuromuscular injuries in the last 6 months, and orthopedic, neurological, cardiac, vestibular, or severe visual impairments. Subjects who were unable to initiate gait without a walking aid or took medication that negatively affected balance or gait were excluded from the experiment. The experimental protocol was approved by the ethical committee of UZ Leuven (number S64605) and subjects signed informed consent.

**Table 1.** Descriptive statistics of study population.

|  | Old | Young | p-value |
| --- | --- | --- | --- |
| Age [Median [min, max] (years)] | 70 (67,89) | 22 (20,29) | - |
| MOCA score [Median (min, max)] | 28 (24,30) | 29 (25,30) | 0.07 |
| FES-I score [Median (min, max)] | 17 (16,35) | 17 (16,22) | 0.38 |
| Gender [Female (%)] | 13 (52) | 12 (48) | 1.00 |
| Leg dominance [Right dominant (%)] | 16 (64) | 23 (92) | 0.04 |

Notes: Age, Montreal cognitive assessment (MOCA) score, and Fall efficacy scale international (FES-I) score, gender, and leg dominance for our study sample. P-values are based on nonparametric tests for MOCA score and FES-I. Between group differences in gender and leg dominance were assessed using chi-square test.

### Protocol

#### Step initiation experiment

Subjects performed a stepping task in which they stepped into projected targets starting from quiet standing (Figure 1B). Targets were constructed in D-Flow (Motekforce link, Amsterdam, Netherlands) and projected on an instrumented split belt treadmill that did not move during the experiment (M-Gait, Motekforce link, Amsterdam, Netherlands). Target size and location were determined based on the subject’s body dimensions. To assess performance trade-offs, accuracy, speed, and stability requirements were altered by manipulating target location and size or altering the instructions in four conditions.

**Figure 1.**
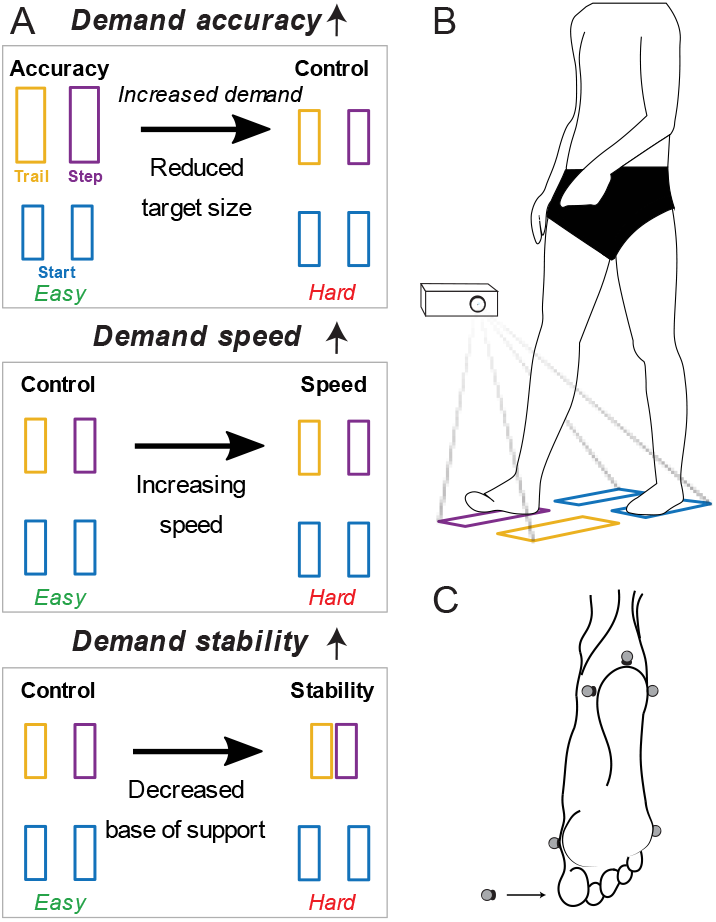
Graphical impression of the experimental protocol. The effect of increasing accuracy, speed, and stability demands on stepping was tested by assessing the change in stepping outcomes as a result of reducing target size, increasing speed, and decreasing the base of support (panel A top to bottom). The horizontal arrows in panel A indicate in which conditions demand was increased. Subjects were asked to step into targets projected from standstill (panel B). In panel A and B, starting targets (blue), stepping leg target (purple), and the trailing leg target (yellow) are positioned as for a right dominant subject with. Panels C shows marker placement to determine positioning of the foot with respect to the targets. Markers were placed on the posterior apex, lateral and medial aspects of the heel, head of metatarsal 1 and 5, and on the nail of the first toe (indicated by the arrow).

- *Control condition*. This condition served as the reference condition. Subjects were asked to step at comfortable pace into rectangular targets with width and length equal to the width and length of the subject’s foot. Step width and length were set equal to spina iliaca anterior superior distance (pelvic width) and 0.5 times the distance between lateral malleolus and trochanter major (i.e. leg length), respectively.
- *Accuracy condition*. Demand on accuracy was reduced with respect to the control condition by increasing the target size by 2cm in width and 5.1cm in length.
- *Speed condition*. Demand on speed was increased with respect to the control condition by instructing the subject to step as fast as possible.
- *Stability condition*. Demand on stability was increased with respect to the control condition by reducing the base of support at the end of the step by reducing step width to 0.5 times pelvic width.

Subjects were instructed 1) to distribute their weight equally over their two feet before stepping to prevent anticipatory weight shifts, 2) to step into the targets with the dominant side first as soon as the targets appeared, 3) not to reposition their feet after placing the feet in the targets, 4) to step as accurate as possible at a comfortable stepping speed except for when explicitly being told to step as fast as possible (speed condition), and 5) to try to stand still as quickly as possible after finalizing the step.

Before the measurement started, subjects performed each condition one time to get acquainted with the experimental procedure. The order of the control, accuracy, speed, and stability conditions was randomized during the measurements.

#### Sensorimotor function

We collected measures of motor variability, proprioception, and reliance on sensory information during standing.

Motor variability was expressed as the variability in torque during a torque matching task (35). Dominant knee and ankle torque was recorded by a Biodex device at 1kHz (Biodex Medical systems, Shirley NY, US). The knee was tested while subjects were seated in upright position with the knee 90° flexed and the ankle slightly plantar flexed. The ankle was tested with the back rest slightly tilted backwards (85°), with upper leg support flexing the hip in 45° and the ankle in neutral position (supplement eFigure 3). The joint’s center of rotation was aligned with that of the Biodex arm. Subjects were asked to perform three maximum voluntary contractions (MVC). The peak moment over these three trials represented MVC in following conditions. Consequently, subjects were asked to produce a torque of 15% MVC, 20% MVC, and 30Nm for 15s guided by visual feedback on a computer monitor. A blue line represented the target torque, a red line represented the produced torque, and a yellow block marked the 15s-time window. Each condition was repeated three times with at least 30s rest between trials.

Knee sensory function was assessed using a position matching task. Subject were seated (approximately 90° knee flexion and 20 degrees ankle plantar flexion) during the test. The examiner imposed target joint angles by passively moving the non-dominant knee. Subjects were then asked to match the examiner-imposed angle with the dominant leg and to indicate the moment at which both the dominant and non-dominant joint angles matched by pressing a hand-held trigger that sent a pulse to the recording system. Target knee joint angles were 10° or 20° flexion. Each target angle was measured 10 times. Joint angles were computed from marker trajectories recorded using motion capture (see below).

Contributions of vestibular, visual, and proprioceptive information to maintain upright posture was assessed by the Sensory Organization Test on the NeuroCom Smart Balance Master (NeuroCom International, Inc, Clackamas, OR, US). Before testing, the subject was positioned on the Balance Master in agreement with device instructions and asked to stand as still as possible. Consequently, postural sway was measured in six different conditions while standing upright: 1) unperturbed, 2) eyes closed, 3) sway-referenced movement of the visual environment (i.e. visual environment moved along with subject such that visual information was reduced), 4) postural sway-referenced movement of the platform (i.e. platform moved such that the subject’s ankle angle remained quasi constant and proprioceptive information was reduced), 5) eyes closed with postural sway-referenced movement of the platform, and 6) postural sway-referenced movement of the visual environment and the platform. Every condition consisted of three 20s trials.

#### Collection of kinetic and kinematic data

During the stepping and joint position matching tasks, thirteen Vicon cameras (Vicon, Oxford Metrics, Oxford, England) measured the trajectories of markers placed on upper legs, lower legs, and feet (supplement eFigure 1) at 100Hz. Six markers on each foot registered foot placement with respect to the targets, see Figure 1C. Ground reaction forces under both legs were sampled at 1kHz by the instrumented treadmill.

## Data analysis

### Kinetic and kinematic data

Markers trajectories and ground reaction forces were processed in MATLAB (R2021a, MathWorks, Natick, USA). After labelling marker data in Vicon’s Nexus 2.11, periods of missing marker data shorter than one second were interpolated. Ground reaction force and marker data were filtered using a 3^rd^ order low pass bidirectional Butterworth filter with a 20Hz cut-off frequency.

Kinematics were estimated using OpenSim 4.1 (36). First, a generic model (gait2392) was scaled to the subject’s dimensions based on a static trial using OpenSim’s Scaling tool. Then, joint angle trajectories for the proprioception trials were estimated using OpenSim’s Inverse Kinematics tool.

### Stepping outcomes

We used foot placement error to quantify stepping accuracy, step duration to quantify stepping speed, and mediolateral center of pressure path length (COPpath_ml_) to quantify stepping stability.

#### Foot placement error

When all six markers on the feet were inside the target, foot placement error was 0. Otherwise, we computed the maximal distance between the foot markers and the target edge in the mediolateral and anteroposterior direction (Figure 1C). Foot placement error was then calculated as the Euclidean norm of the mediolateral and anteroposterior foot placement errors and averaged over both legs. Foot placement error was evaluated over the foot flat phase, i.e. trailing limb toe-off to foot landing for the stepping leg and trailing limb foot landing to trailing foot weight acceptance for the trailing leg. For trailing foot weight acceptance, see the last paragraph of this section.

#### Step duration

The duration of a step was computed as the time between target appearance and trailing foot weight acceptance.

#### Mediolateral center of pressure path length

COPpath_ml_ was calculated over the 1s period after non-dominant (trailing) leg foot landing. The COPpath_ml_ was normalized to pelvic width to be able to compare between individuals. COPpath_ml_ quantified the subject’s ability to stabilize posture after the step according to the instruction to stand still as quickly as possible after finalizing the step.

Stepping events were determined based on ground reaction force data. A threshold of 30N on the vertical component of the ground reaction force was used to detect toe-off and foot landing events. Finally, we defined trailing foot weight acceptance as the moment the vertical ground reaction force increased over 10% of the subject’s body weight in the period after non-dominant foot landing. To represent stepping outcomes for each condition, we calculated the median over five repetitions per condition.

### Trade-offs between task-level goals

We computed performance trade-offs between task-level goals by calculating the difference in stepping outcomes between the condition in which we altered stepping requirements and the control condition. For example, we evaluated the change in step duration (T) between the speed and control condition to evaluate the effect of increasing demand, see Figure 1A second row.

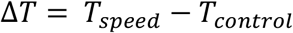

Next, we tested the effect of increasing speed on accuracy and stability by calculating the change in foot placement error (ϵ) and *CoPpath*_*ml*_between the speed and control conditions:

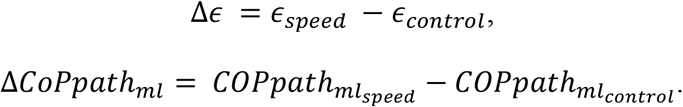

*ΔFoot placement error* and *ΔCOPpath*_*ml*_ represent the speed-accuracy and speed-stability trade-offs respectively. We similarly evaluated the effect of increasing stability and accuracy demands.

### Sensorimotor function outcomes

#### Torque variability

Torques measured from the Biodex were interpolated for missing values after which the signal was filtered using a 5^th^ order bidirectional Butterworth filter with a cutoff frequency of 25Hz. Consequently, torque variability was estimated as the coefficient of variance over a 5s time window, between 5s and 10s after the start of the trial, for each joint and torque level separately. Per condition, torque variability for the ankle and knee was represented as the median coefficient of variance over trials. In the subsequent statistical analysis, we averaged the median coefficient of variance over conditions to obtain one value for the ankle and knee.

#### Proprioceptive accuracy and precision

Joint angle time series were averaged over the 0.1s period after the trigger pulse was recorded. The joint matching error between the two knees was then calculated as the difference between the imposed and matched angle per trial. Finally, the mean and standard deviation in joint matching error over 20 trials were calculated to represent proprioceptive accuracy and precision.

#### Somatosensory (SOM), visual (VIS), vestibular (VEST), and visual preference (PREF) scores

For each of the six sensory organization test conditions, an equilibrium score EQ_i_ with i referring to the condition was computed. Equilibrium scores quantify postural stability by comparing the measured sway against the theoretical sway limit of 12.5° (37). These equilibrium scores were combined to obtain four outcomes describing the subject’s reliance on sensory modalities for postural stability (38): 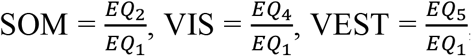, and 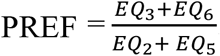 Where the VIS score measures the ability to use visual information to maintain quite standing and the PREF score measures the ability to suppress visual information that is incongruent with the body’s spatial orientation.

### Statistical analysis

To determine whether trade-offs between task-level goals in stepping were larger in older compared to young adults, the six trade-off measures were compared between age groups using Multivariable Analysis of Variance (MANOVA). We considered stepping performance trade-offs as separate variables in the MANOVA. Consequently, we identified differences between age groups by means of two-tailed pairwise comparison. We corrected for the effect of multiple comparisons based on the Benjamini-Hochberg procedure. Test outcomes were considered significant below an alpha value of 0.05.

Finally, we explored the hypothesis that age-related differences in task-level goal trade-offs were associated with decreases in sensorimotor function. Before calculating correlations, data was mean corrected for each group. When statistical assumptions were met, we tested relationships using Pearson correlations. Otherwise, we used Spearman’s rho (*p*_*spearman*_) to assess relationships. As in this part of the analysis we tested our exploratory hypothesis, we corrected correlations for multiple comparisons for proprioception, torque fluctuation, and sensory organization test outcomes separately. One older subject was removed from the analysis of proprioceptive and force fluctuation data as this subject was unable to comply with the measurement protocol.

The statistical analysis was carried out in R (R Foundation for Statistical Computing, Vienna, Austria), version 4.1.3 (39).

#### Preregistration

Hypotheses, the experimental protocol, data processing, and analysis steps were preregistered on the Open Science Framework website (40). Deviations from the preregistered documents are described in the supplementary material.

## Results

### Preregistered outcomes

We found trade-offs between task-level goals as we increased stepping demands (V = 0.35, F(6,43) = 3.9, p < 0.01) but these were not different between young and older adults. Foot placement error increased with increasing demand on speed and stability (median = 0.43 cm, interquartile range (iqr) = 1.25 cm, p = 0.03 and median = 0.47 cm, iqr = 0.88 cm, p < 0.01 respectively, Figure 2 panel B and C). MANOVA test results indicated that trade-offs were different between young and older adults (V = 0.25, F(6,43) = 2.41, p = 0.04). However, after performing pairwise tests with correction for multiple comparisons, we did not find a difference between groups.

**Figure 2.**
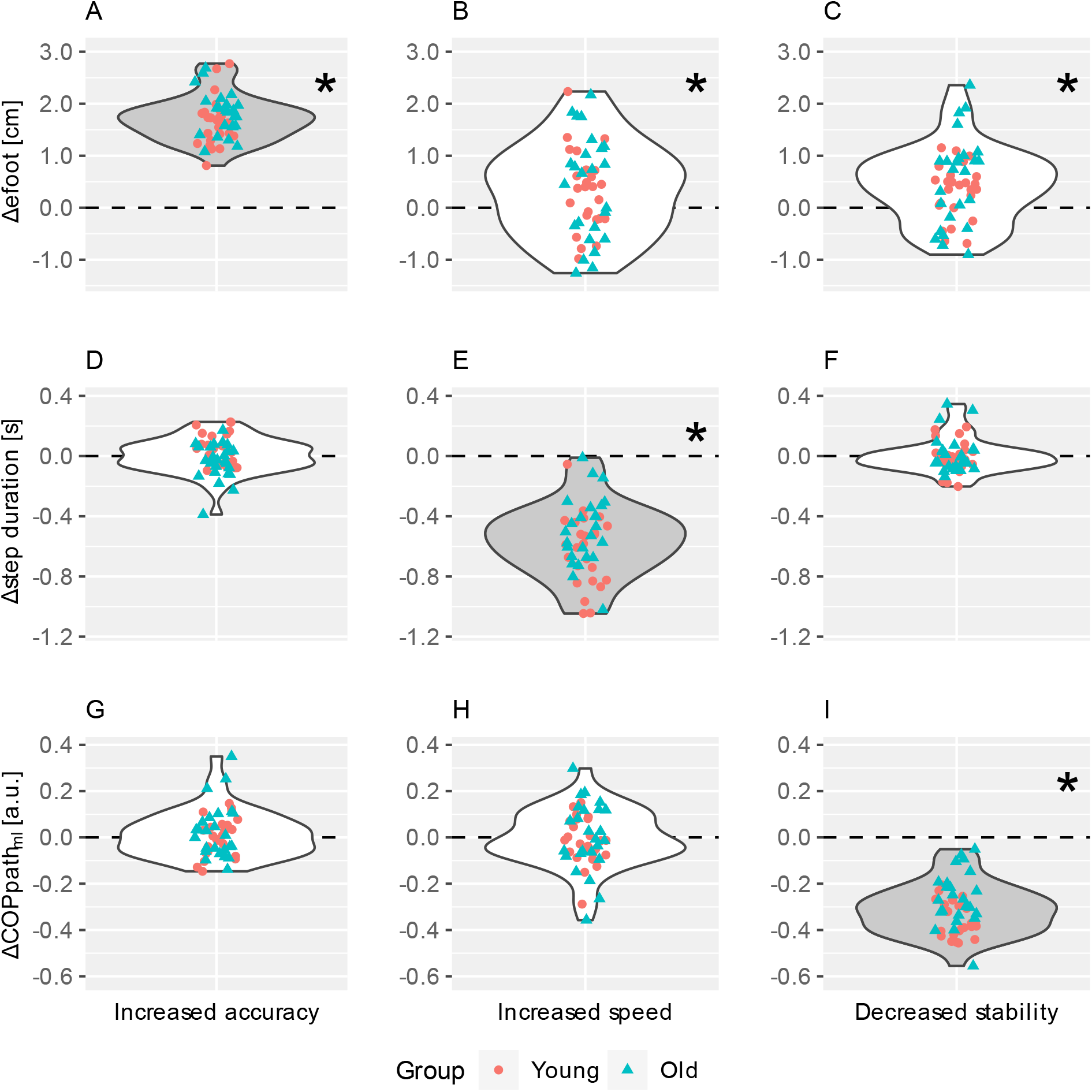
Stepping outcomes with increasing task-demands. Circles and triangles represent individual data points for young and older adults respectively. Plots show the change in foot placement error, step duration, and mediolateral center of pressure path (in the rows in top-to-bottom order) as a result of increasing accuracy, speed, and stability demands (in the columns in left-to-right order). Panels A, E, and I, show changes in task-level goal performance (i.e. stepping outcomes) on task-goals in which demand was increased. These plots thus show the effectiveness of increasing task demands. E.g. panel E shows whether increasing demand on speed successfully decreased step duration. Panels B, C, D, F, G, and H show the change in stepping performance resulting from task-level goal trade-offs, i.e. the extent to which increased demand on one task goal leads to decreased performance on the others. E.g. panel B illustrates the change in foot placement error resulting from increasing speed, i.e. the speed-accuracy trade-off. *Indicates when increasing stepping demands resulted in a change in stepping outcomes that was statistically significant different from zero after correcting for multiple comparisons.

**Figure 3.**
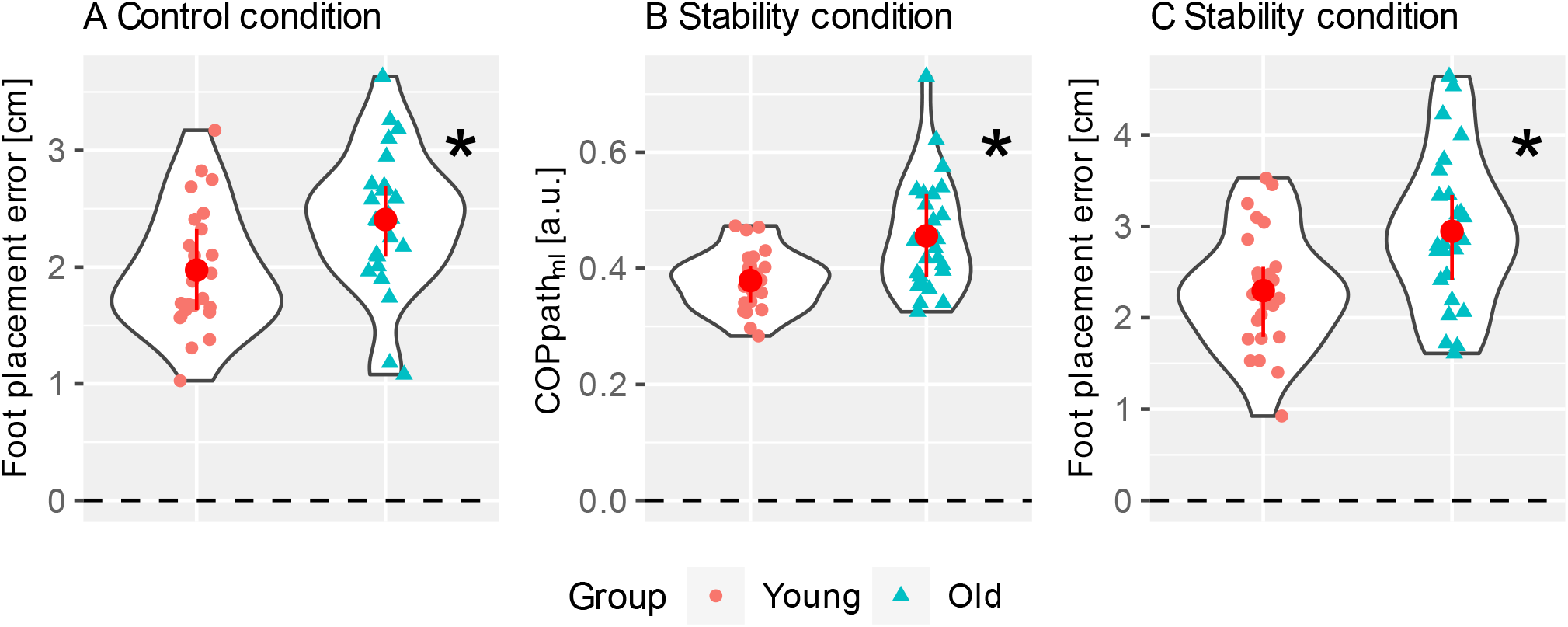
Significant differences in stepping outcomes between young and older adults. Between group differences in foot placement error in the control condition (Panel A), and mediolateral center of pressure path and foot placement error in the stability condition (Panel B and C) with individual data points for young (circle) and older adults (triangles). The group median and interquartile range are represented by the red circle and whiskers.

We found few correlations between task-level goal trade-offs and sensorimotor function. Changes in mediolateral COP path length with increasing demand on speed correlated with SOM score (*p*_*spearman*_= 0.47, p = 0.01). Correlations between trade-offs and other sensorimotor function outcomes (i.e., knee and ankle torque variability, knee joint matching error and standard deviation matching error, and other sensory ratios) did not reach significance after correction for multiple comparison.

### Unregistered outcomes

The accuracy, speed and stability conditions successfully increased demand on stepping requirements (V = 0.97, F(3,46) = 575.06, p < 0.001). Foot placement errors increased when target sizes decreased (median = 1.73 cm, iqr = 0.49 cm, p < 0.001), step duration decreased when subjects were instructed to step faster (median = −0.55 s, iqr = 0.30 s, p < 0.001), and mediolateral COP path decreased when step width in the target position decreased (median = − 0.31 a.u., iqr = 0.14 a.u., p < 0.001), Figure 2 panel A, E, and I. These results indicate that stepping conditions successfully challenged accuracy, speed, and stability.

Stepping performance was poorer in older than in young adults. Foot placement error was larger in older compared to young individuals in the control and stability condition. In the control condition, median foot placement error was 2.41 cm (iqr = 0.61 cm) in older and 1.88 cm (iqr = 0.69 cm) in younger adults, p = 0.03. In the stability condition, median foot placement error was 2.80 cm (iqr = 0.93 cm) in older and 2.26 cm (iqr = 0.77 cm) in younger adults, p = 0.03. Also in the stability condition, mediolateral COP path length was larger in older (median = 0.43 a.u., iqr = 0.14 a.u.) compared to young adults (median = 0.38 a.u., iqr = 0.06 a.u.), p = 0.03.

Knee joint position matching error and the standard deviation of the matching error were not different between young and older adults (Table 2). Additionally, the coefficients of variance in torque production for the knee and the ankle were not different between young and older adults (Table 2). Two of the four scores on the sensory organization test were lower in older adults. The VIS score was lower in older (median = 0.80) compared to young adults (median = 0.86), p < 0.01, indicating that reducing proprioceptive information induced a larger increase in sway in older than in young adults. VEST score was also lower in older (median = 0.57) compared to young adults (median = 0.72), p < 0.01, indicating that reducing proprioceptive and visual information induced a larger increase in sway in older than in young adults.

**Table 2.**
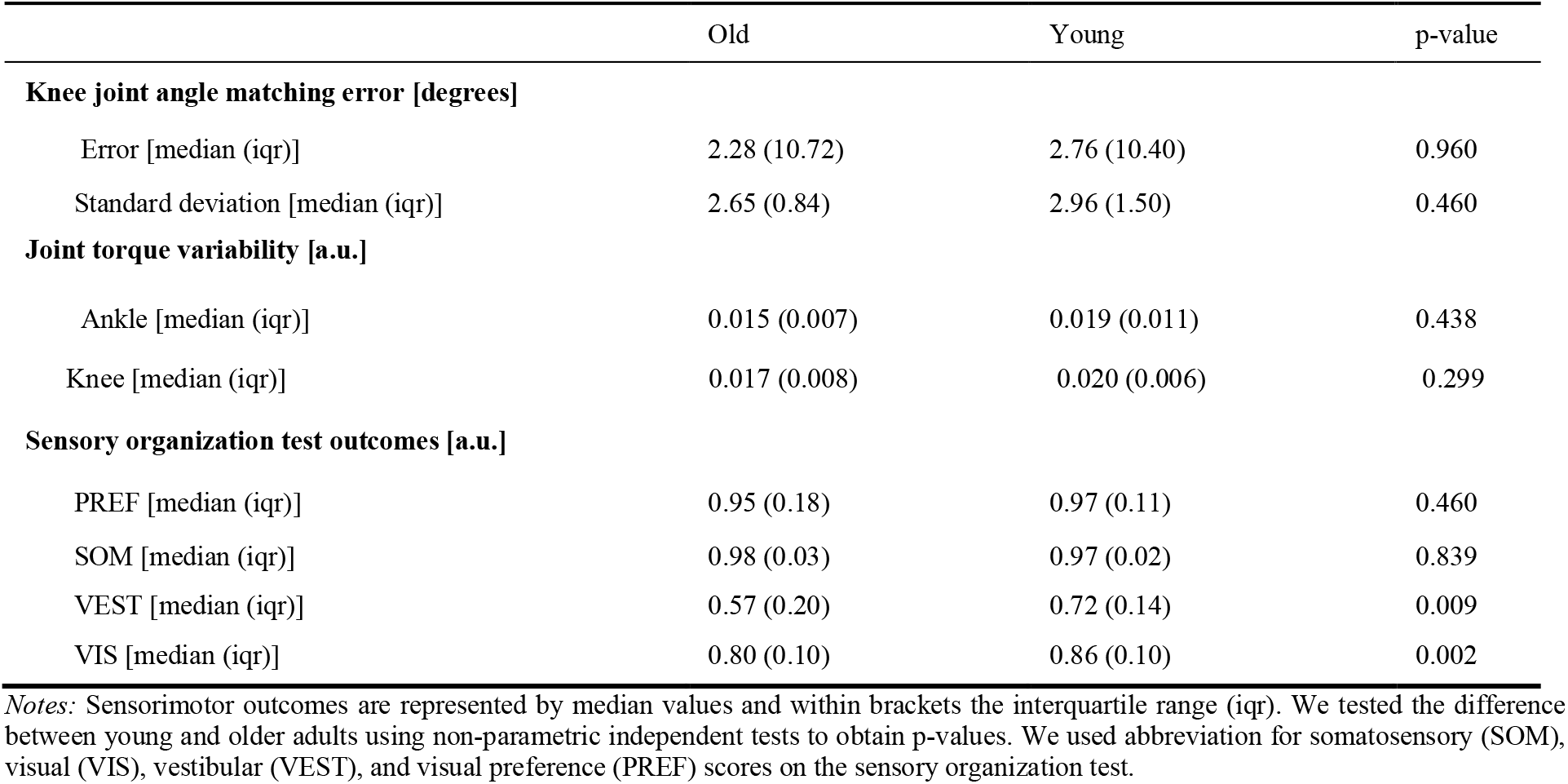
Between group differences in sensorimotor outcomes

Raw data and processing scripts were published for replication and reuse (41).

## Discussion

We found no differences in the magnitude of the accuracy-speed-stability trade-off between young and older adults although stepping performance was worse in older adults. Older adults stepped less accurate than young adults in the control and stability condition, indicating clear differences between groups. In the stability condition, older adults also showed more center of pressure movements at the end of the step, suggesting that they needed more balance correcting movements than young adults when stability requirements were high. As we did not find differences in accuracy-speed-stability trade-offs between groups, reduced stepping accuracy and stability in older adults was not caused by a reduced ability to combine different task-level goals in the stepping task. We thus reject our confirmatory hypothesis that larger task-level goal trade-offs contribute to decreased performance in older adults. Additionally, sensorimotor function could not explain inter-individual differences in task-level goal trade-offs. We therefore also reject the hypothesis that poorer sensorimotor function is related to larger trade-offs between stepping accuracy, speed, and stability. Nevertheless, we found that older adults have similar speed-accuracy and accuracy-stability trade-offs to young adults. Due to the accuracy-stability trade-off, a reduced ability to control balance may have led to less accurate steps in older adults.

### The observed speed-accuracy trade-off across age groups is partly in line with previous findings

We observed that foot placement error increased when asking participants to step as fast as possible compared to stepping at self-selected speed. Reynolds and Day (20) found a similar increase in foot placement errors when giving young adults 300ms (fast) instead of 600ms (slow) to complete their step. However, our observation, that stepping speed did not decrease when increasing accuracy demands, contrasts a previous study by Drury and Woolley (11). They found a decrease in speed when reducing target size. These contrasting findings may be due to differences in experimental conditions. First, the difference in stepping accuracy requirements between conditions was smaller in our study (approx. 20%) than in the study of Drury and Woolley (approx. 300%). Second, we imposed stepping accuracy less stringently than Drury and Woolley. Drury and Woolley requested subjects to repeat the step in case of a foot placement error (11).

### The observed accuracy-stability trade-off across age groups suggests a conflict between stepping to the target and foot placement to control balance

Similar to our findings, Nonnekes and colleagues (22) found a decrease in stepping accuracy in stroke survivors and age matched controls when the base of support at the end of the step was decreased in a stepping task. This indicates that reducing the base of support at the end of the step causes a conflict between stepping to the target and foot placement to control balance. In the same study, Nonnekes and colleagues found that stepping accuracy increased when individuals, and especially stroke survivors, used crutches that increased their base of support during the stepping task. This effect is also seen in walking where the use of crutches improved foot placement accuracy in both young and older adults (23). Hence, conditions that improve balance, either through a wider step as in our study or upper limb support as reported in literature, improve stepping accuracy, probably by reducing the need for foot placement as a balance control strategy.

### Reduced ability to control balance may have led to less accurate steps in older adults

Several observations seem to indicate that our group of older adults had a reduced ability to control balance. First, mediolateral COP path length was larger in older adults when the base of support at the end of the step was decreased with respect to the control condition. Second, older adults scored lower on the sensory organization test conditions in which the platform was sway referenced, i.e. a lower score on EQ4, EQ5, and EQ6 (supplement eTable 1). Given the accuracy-stability trade-off (in both young and older adults), lower stability in older than young adults may have been the origin of less accurate stepping in the stability condition. Possibly, reduced accuracy in the control condition may be attributed to reduced stability. We did not observe any differences in stability in the control condition between young and older adults based on COP path length at the end of the step, but our stability measure might not have captured all age-related differences in stability. In general, it is hard to determine stability in experiments that did not cause a loss of balance (42).

### Older adults had the same preferred stepping speed as young adults, which is in contrast to previous observations

For example, older adults have repeatedly been observed to step slower than young adults in a choice stepping task, in which subjects step to targets that appear in multiple locations (8,13,43,44). In these earlier studies, authors hypothesized that older adults increase the time in the weight shifting phase because they prioritize stability over stepping speed (8,13). Despite that weight shifting was a prerequisite in this study’s stepping task, older adults were not slower than young adults. We did not find a speed-stability trade-off in both age groups indicating that both young and older adults are able to increase speed without compromising balance. A difference in cognitive load between paradigms may explain the different findings. The cognitive load is higher in a choice stepping task where the target direction is variable in contrast to our stepping task where the target always appeared in front of the subject. Increased cognitive load was previously found to amplify age-related differences in stepping (14).

### Inter-subject variability in trade-offs cannot be explained by variability in sensorimotor function

as we found few associations between sensorimotor outcomes and trade-offs. We explored 48 potential associations between trade-offs and sensorimotor function and found only one positive association between the change in mediolateral COP path length with increasing speed and the somatosensory score on the sensory organization test. Individuals with a low somatosensory score, i.e. low equilibrium score in the eyes closed relative to the eyes open condition, depend more on visual information to stand still. Hence, the association between COP path and somatosensory score would indicate that individuals who relied more on visual information for postural balance reduced mediolateral COP path length more when speed increased. However, it is questionable whether further research could confirm this relationship as we only partially accounted for multiple comparisons, i.e. only within proprioception, torque variability, and sensory organization test outcomes separately. Since we only find one relationship among the many relationships we tested, we conclude that variability in trade-offs were poorly explained by sensorimotor outcomes.

### Task-level goal trade-offs and sensorimotor deficits may be larger in fall-prone compared to healthy older adults

In stroke survivors, increasing the base of support resulted in larger improvements in stepping accuracy than in age matched healthy controls (22). This suggests that accuracy-stability trade-offs may be larger in individuals with balance deficits. Moreover, it has been suggested that the lower stepping speed in fall prone older adults compared to healthy older adults may be due to a larger stability-speed trade-off in fall-prone older individuals (45). Yet, this remains to be investigated. If accuracy-speed-stability tradeoffs are indeed larger in fall prone compared to healthy older adults, testing for relationships between sensorimotor function and trade-offs may shed light on potential mechanisms. Sensorimotor function, and in particular proprioception, was found to predict falls (33,46) indicating that the relationship between stepping stability and sensorimotor function may be stronger in fall-prone compared to healthy older adults.

## Acknowledgments

We would like to thank all subjects for participating in the experiments. During the experiments we received support from master students. In particular, we would like to thank Dries De Moor and Michiel Follon for their support during the experiments. We received support from Steffen Fieuws (Leuven Biostatistics and Statistical Bioinformatics Centre) on the design of the statistical analysis. The authors would like to thank Jente Willaert for reading the manuscript and providing feedback.

## Funding

This work was supported by ‘Fonds Wetenschappelijk onderzoek’ (FWO), Research foundation Flanders. The project was funded by a FWO research project (G088420) received by FDG and a FWO fellowship (1SF7322N) received by WM.

## Conflict of interest

None declared.

## Author Contributions

F.D.G., M.A., and W.M. developed the study concept. F.D.G., M.A., S.A., C.D., and W.M. designed the experiments. After the design of the study F.D.G, M.A., and W.M. preregistered the study. S.A., C.D., and W.M. acquired the data. W.M. processed and analyzed the data. F.D.G., M.A., and W.M. interpreted the results, wrote, and edited the manuscript.

## Notes

https://doi.org/10.48804/6GRWWE

### Competing Interest Statement

The authors have declared no competing interest.

