## Supplementary material for "Accuracy-speed-stability trade-offs in a targeted stepping task are similar in young and older adults"

#### Estimation stepping target location

Targets were projected on the treadmill using a projector that was bolted to the floor to prevent movement of the projector relative to the projection surface. The projection was constructed using DFlow software. To project targets scaled to the dimensions of the subject, we converted body dimensions from meters to DFlow units using separate scaling factors for the mediolateral and anteroposterior dimensions. The scaling factors were calculated by dividing the width and length of the projection surface in DFlow units by the width and length measured in meters using marker positions. Target location, width, and length were saved in DFlow units for the different conditions. The targets were projected under a slight angle with respect to the projection surface (and thus marker coordinate frame) due to the low resolution by which the projection could be rotated in DFlow. Target location, width, and length saved in DFlow did not account for this rotation with respect to the projection surface. As we relied on reconstruction accuracy of the targets for computing foot placement error, we calculated the rotation between the targets and the marker coordinate frame. Consequently, we estimated the reconstruction accuracy of targets by assessing the error between measured marker locations on the projection surface and their estimated location based on DFlow data using calculated scaling factors and the rotation. The mean error between the estimated and measured marker positions was 3.4mm. The reconstruction errors were larger towards the edges of the treadmill, possibly because of nonlinear distortion of the projected image. Nevertheless, this will have minimally impacted the estimated target locations as targets were projected in the middle of the treadmill where the errors were relatively small.

#### Correction error Vicon camera calibration

Due to an error during calibration of the Vicon cameras, for two of the subjects (subject 40 and 41) the marker coordinate frame was translated with the respect to the force plate coordinate frame. We corrected the error in the processing script by translating the marker coordinate frame to match the force place coordinate frame.

#### Protocol marker placement

We measured markers on the upper legs, lower legs, and feet to register the position of leg segments during the proprioception and stepping trials (eFigure 1). To measure the position of the upper leg we used a 3-marker cluster together with markers on greater trochanter, medial, and lateral epicondyle of the thigh bone. We measured the position of the lower limb segment using a 3-marker cluster and markers on the medial and lateral malleolus. The position of the foot was measured using markers on the calcaneus' posterior, medial, and lateral aspects, the bases of metatarsal 1 and 5, and the nail of the big toe.

### eFigure 1. Marker positions

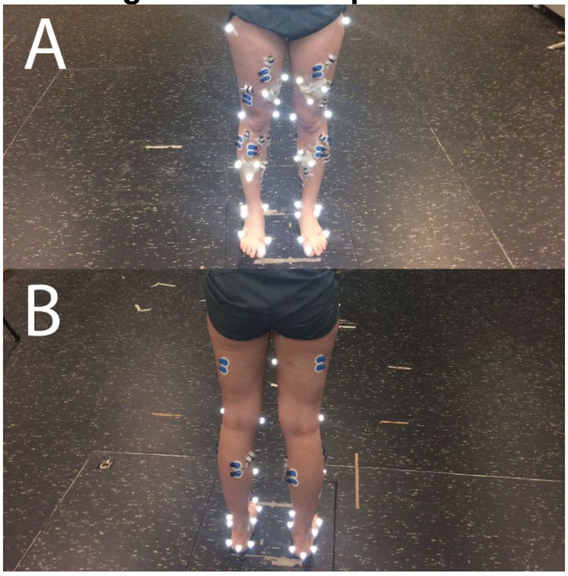

Impression of marker positions on the anterior (A) and posterior (B) sides of the lower limbs. Besides markers, we also placed EMG electrodes on lower limb muscles. The analysis of the EMG data is out of the scope of this paper.

**Assessing proprioception**

In addition to procedures described in the methods of the paper, we instructed subjects to actively extend their non-dominant knee against resistance for approximately 3s to reduce the influence of thixotropy before each proprioception trial (1). Besides the knee angle position matching trials, we estimated joint angles in a reference posture in which both legs were aligned. We measured marker position in this reference posture calculated knee joint angles using OpenSim's Inverse Kinematics Tool. Consequently, we computed differences between joint angles of the dominant and non-dominant leg which were due to inconsistencies in marker placement and tissue artifacts. We corrected proprioception trials for this error in knee joint angles.

**Weight distribution task in stepping trials**

To help subject in equal distribution of body weight over their two legs, subjects were informed about how they distributed their weight over both feet. Two blocks, blue and red, were projected on the treadmill in front of the subject. The anteroposterior position of the blocks was proportional to the ground reaction force under the left (blue) and right (red) leg (eFigure 2). Subjects were asked to align the two blocks by redistributing weight over their legs. Weight distribution had to be within 40% to 60% bodyweight for each leg for five seconds before the stepping targets were presented.

### eFigure 2. Weight distribution task

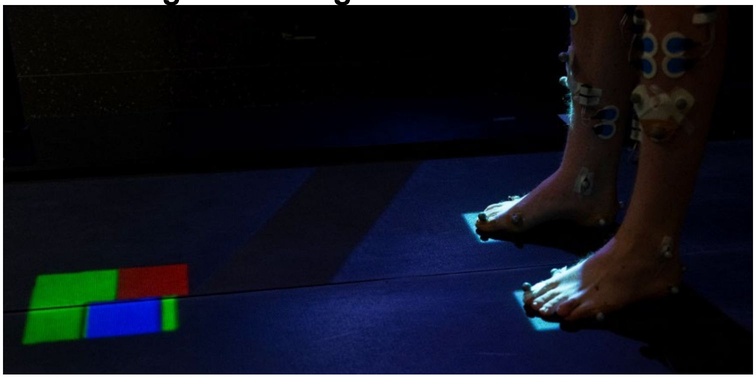

Subjects performed a weight distribution task before stepping targets were presented. The anteroposterior position of the blue and red blocks was proportional to the ground reaction force under the left (blue) and right (red) leg. The targets would only appear after the subject matched the position of the red and blue block for 5 seconds, by positioning them into the green area. We allowed for a difference of 20% body weight between the right and the left leg.

### eFigure 3. Impression torque fluctuation and proprioception tests

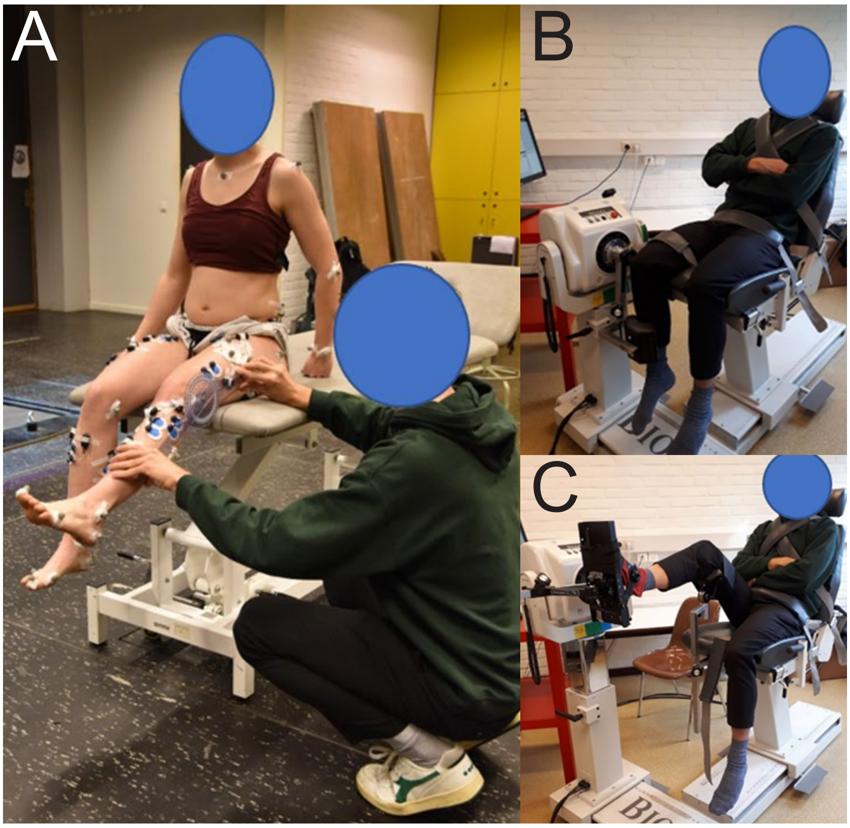

Impression of body configurations during proprioception (A) and ankle (B) and knee (C) torque fluctuation tests. Panel A, the examiner imposed target joint angles by passively moving the non-dominant knee while the subject was seated (approximately 90° knee flexion and 20° ankle plantar flexion). Panel B, the knee was tested while subjects were seated in upright position with the knee 90° flexed. Panel C, the ankle was tested in a similar body configuration, but with the back rest slightly tilted backwards (85°). The ankle of the dominant leg was placed in neutral position by resting the foot on a footplate attached to the Biodex arm, with dominant hip flexed by supporting the upper leg using a leg rest.

**Deviations from preregistration**

*We registered to included subjects scoring 26 point or higher on the MoCA (Montreal Cognitive Assessment).* After data collection we noted that some individuals scored lower than 26 points on the MoCA in both young and older adults. As Table 1 of the article indicates, we did not find a difference between age groups on the MoCA score. For this reason, in combination with that we experienced difficulties in including older adults when COVID measures were in place, we decided to include all subjects into the analysis.

*We changed the instable stepping conditions.* In the preregistered document we described to use tandem stance for the unstable conditions. To prevent overlap between the targets marking the starting location and the stepping targets in the tandem stance condition, we shifted the location of the stepping targets forward. This forward shift resulted in a step that required a larger forward displacement of the center of mass (COM). This meant that step duration was not expected to be equal between the stability condition and other conditions as the average anteroposterior displacement of the COM was larger in the tandem stance. This was undesirable since in the stability condition we intended to test the effect of instable posture and not of COM displacement. Departing from our preregistered document, we decided to use a stability condition in which the base of support was decreased but without a change in mean anteroposterior COM displacement by decreasing the mediolateral distance between the projected targets.

*We did not used planned dimension reduction methods.* We deviated from our plan to use factor analysis to reduce dimensionality of sensorimotor function outcomes. First, we noticed that correlations between sensorimotor outcomes was generally low (<0.6) possibly resulting in low variance accounted by the resultant factors, approximately 70% depending on the approach (not reported). Additionally, factors did reduce the dimensions of the data little more than averaging over sensorimotor outcomes per joint (e.g., for force fluctuation at 30Nm, 15%, and 30% of MVC). Therefore, we decided to reduce the number of outcomes by averaging proprioception and torque fluctuation outcomes over joints, and combining equilibrium scores on the Sensory Organization Test conditions following literature (2). We consider these outcomes to be better interpretable.

*For some (especially) older individuals 5 degrees dorsal flexion was close to the end of the passive range of motion of the ankle.* These individuals may have been able to determine the angle of the ankle based on the force that was applied by the experimenter on the foot. We therefore doubted whether ankle joint position matching errors measured using our protocol were a reliable representative of ankle proprioception. We thus excluded ankle joint proprioception from further analysis.

*We changed the definition of the movement termination event in the paper.* In the paper we defined movement termination as trailing leg weight acceptance, we defined movement termination as the moment at which the pressure under the trailing foot was higher than 30% body weight in the period after trailing toe-off. As this event does not represent the end of the step, we changed the name of this event. Additionally, during the data analysis we noticed that some subjects tended to load their trailing foot less just after the step. As a result, we were less able to reliably estimate step duration since step duration was estimated based on trailing leg weight acceptance. Trailing leg weight acceptance was thus more reliably detected when this percentage was lower, therefore we set this percentage to 10%.

We noted an error in the "Measured variables” section of the preregistration document. In this section we wrote that mediolateral margin of stability is used to quantify stability. This is a remnant of old text and should have been “mediolateral center of pressure path to quantify stability”.

**Assumptions statistical tests**

Before applying parametric tests, we tested associated assumptions. In the case of the MANOVA, we 1) assessed multivariate normality by testing residuals after fitting the MANOVA against the normal distribution, 2) evaluated homogeneity of variance-covariance using the Box’s test, and 3) established linearity by evaluating scatter plots between all pairs of dependent variables. In case of pairwise tests, we tested whether the assumptions of normality and homogeneity of variance were violated when applying independent t-tests in multiple comparisons using the Shapiro-Wilk and the Levene’s test. When assumptions were violations we performed non-parametric tests, i.e. Mann-Whitney U test. Before testing relationships between variables using Pearson moment correlation, we assessed linearity, homoscedasticity, normality, and the existence of outliers.

**Control for speed**

In our experimental setup, stepping speed was not tightly controlled as the comfortable and fast stepping speed may vary among individual subjects. This may have affected the differences we found between young and older adults on stepping outcomes, i.e. in the control and stability conditions. Since, the accuracy-speed trade-off could have affected accuracy, we statistically assessed whether differences in accuracy and stability between age groups were the result of differences in speed. We fitted a linear model between step duration and the other performance outcomes (i.e. foot placement error). Next, we tested for differences between young and older adults by performing statistical tests on the residuals of the linear model. We considered differences in performance between age-groups that prevailed in model residuals not to be the result of differences in self-selected speed due to the accuracy-speed trade-off.

After statistically correcting for stepping speed, older adults still performed worse on the stepping task than young adults. After fitting the linear model, residuals in foot placement error were larger in older (median = 0.27 cm, interquartile range (iqr) = 0.69 cm) compared to young adults (median = -0.31 cm, iqr = 0.77 cm) in the control condition, $p < 0.05$. Likewise, in the stability condition, residuals in foot placement error were larger in older (median = 0.11 cm, iqr = 1.06 cm) than in young adults (median = -0.31 cm, iqr = 0.87 cm), $p < 0.05$. We conclude that age-related differences in step duration did not account for larger foot placement errors in older adults. Between age group differences in equilibrium score

### eTable 1. Between group differences in equilibrium score for each sensory organization test condition

| Conditions | median old | iqr old | median young | iqr young | test statistic | p-value |
|---|---|---|---|---|---|---|
| C1 | 94.67 | 2.33 | 95.33 | 1.67 | 261.00 | 0.40 |
| C2 | 92.00 | 4.33 | 93.00 | 2.33 | 268.50 | 0.40 |
| C3 | 90.67 | 4.33 | 92.67 | 5.00 | 265.50 | 0.40 |
| C4 | 77.00 | 14.00 | 83.67 | 8.33 | 124.50 | 0.00 |
| C5 | 55.33 | 18.67 | 68.67 | 15.00 | 152.00 | 0.01 |
| C6 | 44.67 | 25.00 | 64.00 | 18.33 | 157.00 | 0.01 |

*Note*: The table shows median and interquartile ranges (iqr) for equilibrium scores for each of the six conditions. We tested differences between young and older adults young a non-parametric test and corrected p-values for multiple comparisons. Older adults scored lower on conditions 4, 5, and 6. In these conditions the standing platform is sway referenced, hence in these conditions older adults were less stable. We use this table in the discussion to motivate that stability in older adults was lower compared.
